# Bovine-derived H5N1 influenza virus efficiently infects lactating swine via the mammary gland

**DOI:** 10.64898/2026.07.18.739312

**Authors:** Min Liu, Natalie N. Chillson, Emma A. Martin, Hannah J. Cochran, Janice Y. Park, Brady O’Boyle, Devra Huey, Kara N. Corps, Andrew S. Bowman, Cody J. Warren

**Affiliations:** Department of Veterinary Biosciences, The Ohio State University, Columbus, OH, USA; Infectious Diseases Institute, The Ohio State University, Columbus, OH, USA; Department of Veterinary Preventive Medicine, College of Veterinary Medicine, The Ohio State University, Columbus, OH, USA; Comparative Pathology & Digital Imaging Shared Resource, The Ohio State University Comprehensive Cancer Center, Columbus, OH, USA

## Abstract

Since 2024, highly pathogenic influenza A(H5N1) viruses have spread extensively among U.S. dairy cattle, where they replicate efficiently in the mammary gland and are shed at high titers in milk. To directly assess susceptibility of commercial swine populations to bovine-derived H5N1 virus, lactating sows with prior influenza virus vaccination histories representative of U.S. commercial swine production systems were inoculated via the intramammary route and co-housed with their 1-week-old piglets to evaluate disease outcomes, viral replication, and potential for vertical transmission. Intramammary inoculation of lactating sows resulted in sustained viral RNA shedding in milk, while piglets exhibited sporadic oral viral RNA positivity that mirrored viral kinetics in milk. Lesions in mammary tissue and viral antigen staining, as well as development of neutralizing antibody responses and changes in milk color and consistency, further confirmed infection in the sows. Despite these molecular findings, none of the animals developed overt clinical disease, and respiratory involvement was not noted during the study period. Collectively, we demonstrate that intramammary exposure results in productive influenza A(H5N1) virus infection in lactating sows despite their vaccination histories, indicating the potential threat of viral spillover into commercial swine populations. The clinically inapparent nature of infection presents a risk of subclinical spread and underscores the importance of expanding viral surveillance to swine.

## INTRODUCTION

In recent years, influenza A(H5N1) viruses belonging to clade 2.3.4.4b have spread globally, driving a panzootic that has significantly threatened agricultural production, wildlife populations, and human health. Wild migratory birds introduced influenza A(H5N1) into North America during 2021-2022^1^, resulting in continued outbreaks in commercial and backyard poultry flocks that have, to date, necessitated the culling of more than 200 million birds (USDA APHIS, accessed 07/18/2026)^2^. Wild birds were also implicated in at least four independent introductions of influenza A(H5N1) into U.S. dairy cattle populations that led to widespread transmission among herds, reduced milk production, and substantial economic losses^3–7^. These bovine-adapted H5N1 viruses have additionally been detected in multiple mammalian species, including feral and domestic cats^8,9^, wild carnivores (e.g., foxes and raccoons)^10^, rodents^10^, and humans^11^, underscoring the expanding host range and interspecies transmission potential of contemporary influenza A(H5N1) viruses.

The continued transmission and evolution of influenza A(H5N1) viruses between birds and mammals increases concern regarding the emergence of viruses with pandemic potential. Swine are an important intermediary in this process because they are susceptible to diverse influenza A virus (IAV) lineages and maintain a large pool of endemic IAV genomic diversity. Multiple genetically distinct IAV lineages co-circulate in modern swine production systems due to sustained transmission of endemic strains and new introductions from avian and human reservoirs^12–16^. These conditions provide opportunities for co-infection and genome reassortment, a major driver of rapid viral evolution^17,18^. The emergence of the 2009 H1N1 pandemic virus through reassortment among human, swine, and avian influenza viruses^19,20^ illustrates how swine IAV diversity can give rise to viruses with major public health consequences. Likewise, the establishment of mammalian-adapted influenza A(H5N1) viruses in swine could provide a pathway for further adaptation, reassortment with endemic swine IAV lineages, and emergence of viruses with zoonotic potential.

Beyond their public health relevance, newly emerging IAV strains can have substantial consequences for swine health and production. Within commercial herds, IAV infection is commonly associated with respiratory disease and diminished productivity, and disease may be further exacerbated during co-infection with other respiratory pathogens, increasing morbidity, mortality, and economic losses^21^. High IAV seroprevalence against endemic H1 and H3 IAVs in swine herds, whether through natural circulation or routine vaccination, may lead to the assumption that commercial pigs possess some degree of cross-protection against new virus introductions. However, these immune histories may not fully protect against antigenically distinct viruses such as influenza A(H5N1) and may instead modify or mask disease presentation. This distinction is especially relevant given the recent subclinical detection of H5N1 in a pig in Oregon, demonstrating that spillover into swine can occur^22^. The susceptibility of commercially raised pigs with endemic IAV exposure or vaccination histories to these viruses remains a key knowledge gap^23^.

Prior studies have investigated the susceptibility of swine to influenza A(H5N1) virus infection following respiratory exposure. A recent study by Arruda et al. evaluated the pathogenesis and transmission potential of multiple clade 2.3.4.4b influenza A(H5N1) virus isolates, including viruses of avian and mammalian origin. Although all isolates replicated within the swine respiratory tract, only mammalian-adapted viruses demonstrated efficient replication in the upper respiratory tract and onward transmission^24^. Notably, these infections were largely subclinical, a finding subsequently corroborated by additional studies using bovine-origin H5N1 viruses^25,26^. Together, these studies suggest that swine susceptibility to influenza A(H5N1) following respiratory exposure is limited. Yet the recent emergence of these viruses in dairy cattle has challenged conventional assumptions about IAV tissue tropism. In cattle, the mammary gland is a highly permissive site for influenza A(H5N1) virus replication, with viral RNA and infectious virus detected in milk at levels exceeding those observed in respiratory and systemic tissues^3,27,28^. The pronounced mammary gland tropism of bovine-adapted H5N1 viruses raises the question of whether mammary exposure could similarly support infection in lactating swine.

Here, we used lactating sows with prior IAV vaccination histories representative of commercial swine populations to evaluate susceptibility to bovine-origin H5N1 virus following intramammary exposure. This approach allowed us to directly assess whether these viruses can infect the swine mammary gland under commercially relevant immune conditions. We assessed dose-dependent clinical and virological outcomes, including milk-associated viral shedding, mammary tissue pathology and viral antigen staining, and potential for nursing-associated transmission to suckling piglets. Our findings demonstrate that commercially sourced lactating sows are susceptible to productive mammary infection with bovine-origin influenza A(H5N1) virus when exposure occurs through the teat canal, paralleling the mammary infection phenotype observed in dairy cattle. However, unlike dairy cattle, the clinically inapparent nature of infection in animals with prior IAV immunity may delay detection in the absence of active surveillance. When taken together, these findings identify lactating swine as a potential susceptible host population and underscore the need to incorporate commercial swine into H5N1 surveillance and risk assessment efforts.

## RESULTS

### Clinical observations

To determine the clinical outcome of influenza A(H5N1) in lactating swine, commercial production sows (n = 4) and their respective piglet litters (n = 10 per sow) were acclimated to biosafety level 3 agriculture (BSL-3Ag) containment prior to inoculation (**Fig. 1A**). According to farm records, all sows had received multiple vaccinations against endemic swine influenza A viruses (IAV), consistent with typical commercial practices. We confirmed IAV seropositivity for all sows using hemagglutination inhibition (HAI) and ELISA assays (**Table 1**). Sows were inoculated via the intramammary route with a bovine-origin D1.1 genotype virus (A/bovine/Nevada/WD-210/2025). Genotype D1.1 was selected because it has rapidly expanded across North American wild bird populations^29^, caused multiple independent spillover events into dairy cattle^6,30,31^, and has been associated with severe human infections^32–35^. Animals were exposed to either 1×10^4^ plaque forming units (PFU) (n = 2) or 1×10^6^ PFU (n = 2) of virus per animal, with inoculum distributed equally across half of the actively lactating mammary glands (**Fig. 1B**). These inoculation doses represent typical ranges used in dairy cattle studies^36–40^. Suckling piglets were removed immediately prior to inoculation, returned to their dams at 1-day post-infection (dpi), and co-housed in farrowing crates to allow natural nursing and potential exposure (**Fig. 1C**). One sow and its litter per dose group were euthanized and necropsied at early (3 dpi) and late (14 dpi) timepoints (**Fig. 1A**).

**Figure 1.**
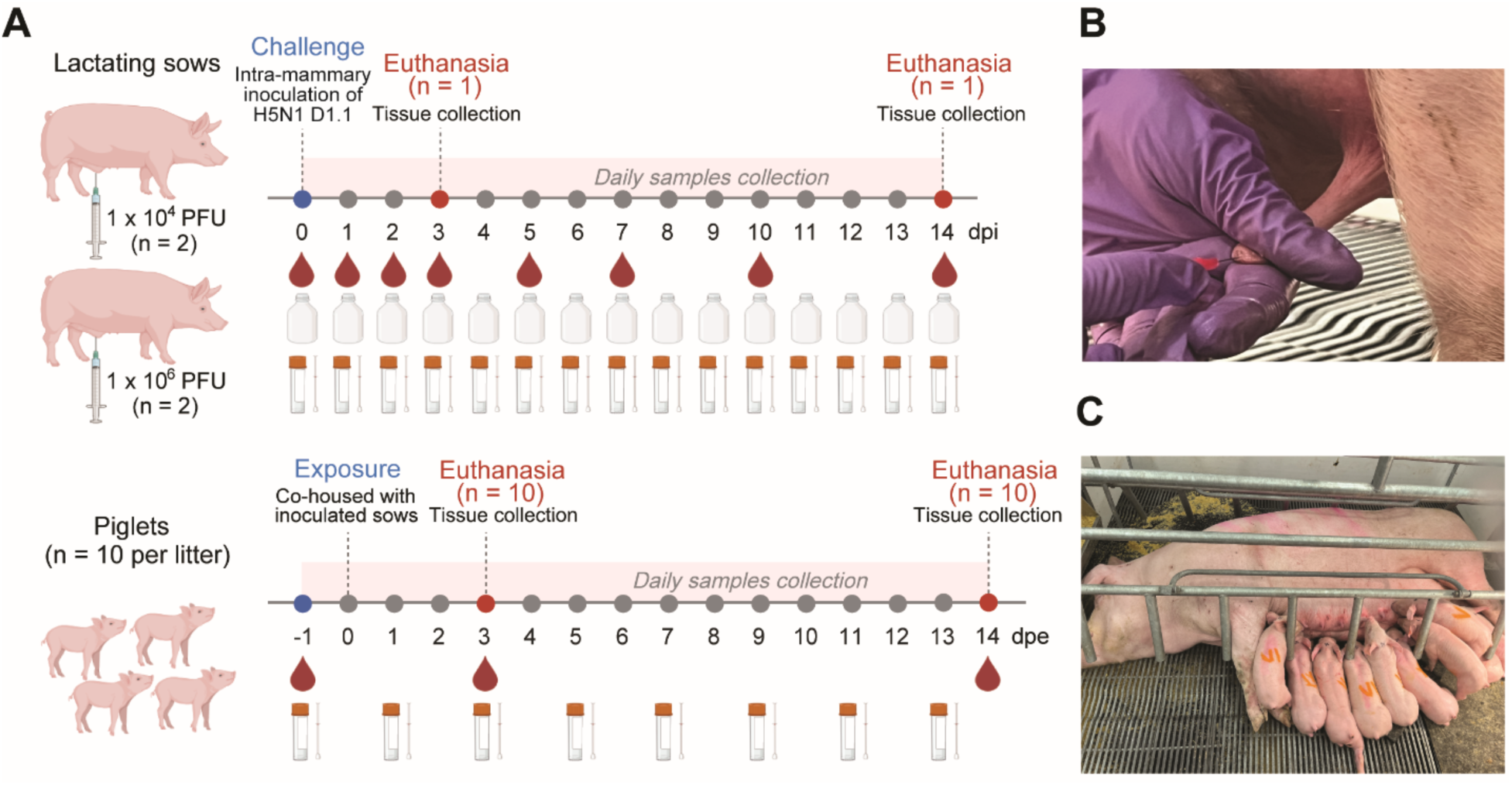
Lactating sow challenge model for assessing influenza A(H5N1) virus infection in swine. (**A**) Schematic of experimental design. Lactating sows were inoculated with 1×10^4^ (n = 2) or 1×10^6^ (n = 2) plaque forming units (PFU) of a bovine-derived influenza A(H5N1) D1.1 genotype virus via the intramammary route. Sows were co-housed with their litters (n = 10 piglets) beginning at 1-day post-inoculation (dpi), which was defined as 0 days post-exposure (dpe) for piglets. Clinical signs were monitored daily, and blood, milk, nasal, oral, and rectal swab samples were collected at indicated time points. One sow per dose group was necropsied at 3 dpi and another at 14 dpi; corresponding piglets were necropsied on the day following the sow’s necropsy. (**B**) Intramammary inoculation was performed by infusing infectious virus into individual teat canals of sows using sterile teat cannula-connected syringes. (**C**) Piglets were reunited with sows 24 h post-inoculation to allow natural nursing and potential viral exposure through suckling.

**Table 1.** Pre-challenge hemagglutination inhibition (HAI) titers and nucleoprotein (NP) ELISA reactivity of swine sera to influenza A virus (IAV)

| Sow ID | <sup>a</sup>HAI assay against diverse IAV subtypes and clades |  |  |  |  |  |  | <sup>b</sup>ELISA |
| --- | --- | --- | --- | --- | --- | --- | --- | --- |
|  | H1 |  |  |  | H3 |  | H5 | IAV NP |
|  | 1A.1.1 | 1A.3.3.3 | 1B.2.2 | 1B.2.1 | 3.2010.1 | 3.1990.4.a | 2.3.4.4b |  |
| Sow#1 | >640 | - | - | - | 320 | 80 | - | 0.54 |
| Sow#2 | >640 | - | - | - | 160 | 40 | - | 0.19 |
| Sow#3 | 320 | - | - | - | - | - | - | 0.64 |
| Sow#4 | >640 | - | - | - | 160 | 40 | - | 0.43 |
| <sup>c</sup>Control gilts | <sup>d</sup>ND | ND | ND | ND | ND | ND | ND | 0.84±0.73 |

<sup>a</sup>HAI titers are expressed as the reciprocal of the highest serum dilution that inhibited hemagglutination.
<sup>b</sup>NP ELISA results are reported as sample-to-negative (S/N) ratios; values >0.67 were considered negative.
<sup>c</sup>Control gilts (n=14) with no prior history of influenza virus vaccination were evaluated via IAV NP ELISA. Data is presented as mean S/N ratio ± SD.
<sup>d</sup>ND, not determined.

Unlike intramammary-inoculation of naïve dairy cattle^36–40^, lactating sows with prior swine IAV vaccination histories did not exhibit overt clinical signs during the study period. Reductions in feed intake at the high-dose condition were transient (**Fig. 2A**), and rectal temperatures remained stable (**Fig. 2B**). Daily observations of piglets identified sporadic cases of diarrhea in both groups (**Supplementary Table 1**); however, no respiratory signs, including sneezing, coughing, or nasal discharge, were observed. Two piglets from the high-dose litter exhibited reduced weight gain and were euthanized early in accordance with pre-established IACUC humane endpoint criteria (**Fig. 2C**). During this period, milk from the high-dose sow (#4) exhibited altered color and consistency (**Supplementary Table 2**), and these changes, potentially in combination with reduced milk output and normal inter-litter competition, may have contributed to the impaired weight gain observed in these piglets. Aside from these cases, piglets appeared clinically normal, maintaining stable weight gain and rectal temperatures over time (**Figs. 2C**, **2D**). Collectively, these findings indicate that commercially vaccinated sows do not develop overt clinical disease following intramammary infection with bovine-derived influenza A(H5N1). Moreover, piglets co-housed with infected sows remained largely clinically unaffected.

**Figure 2.**
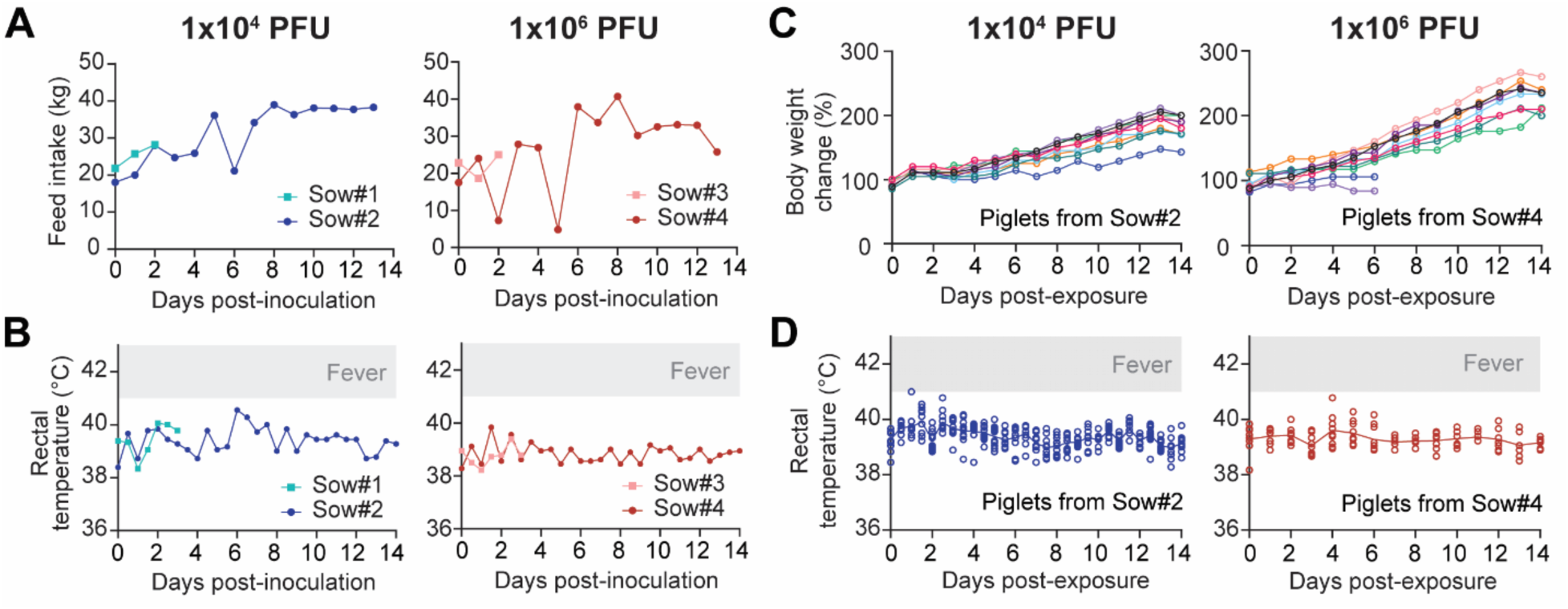
Clinical presentation following influenza A(H5N1) challenge in commercial swine. Commercial lactating sows were challenged via intramammary inoculation with influenza A(H5N1) clade 2.3.4.4b (D1.1 genotype) at doses of 1×10^4^ or 1×10^6^ PFU per sow. At 1-day post-inoculation, suckling piglets (n = 10 per sow) were co-housed with the inoculated sow in farrowing crates. (**A**) Feed intake and (**B**) rectal temperature of sows were monitored daily. Sows #1 and #3 were euthanized at 3 days post-inoculation (dpi), while Sows #2 and #4 were euthanized at 14 dpi. Feed intake represents the total weight of the gruel mixture consumed, consisting of swine dry feed mixed with water at a 1:2 ratio. (**C**) Body weight and (**D**) rectal temperature of piglets were measured daily. Two piglets from Sow #4 exhibited stunted weight gain and were euthanized early based on pre-established IACUC humane endpoint criteria.

### Viral shedding dynamics in lactating sows

Lactating sows possess six to eight pairs of mammary glands; in this study, approximately half of the actively lactating glands were inoculated in a zig-zag pattern (alternating left and right sides), while the remaining glands were left non-inoculated. Visible changes in milk appearance were observed in the high-dose group beginning at 3–4 dpi (**Supplementary Table 2**). Milk from several inoculated glands appeared off-white / yellow and, in some cases, pink-tinged, consistent with possible hemorrhage, whereas milk from non-inoculated glands retained a normal appearance (**Fig. 3A**; representative milk samples from inoculated glands shown on the left and non-inoculated glands on the right). These changes persisted for 5–12 days in affected glands. In contrast, no apparent alterations in milk appearance were observed in the low-dose group (1×10^4^ PFU) (**Supplementary Table 2**).

**Figure 3.**
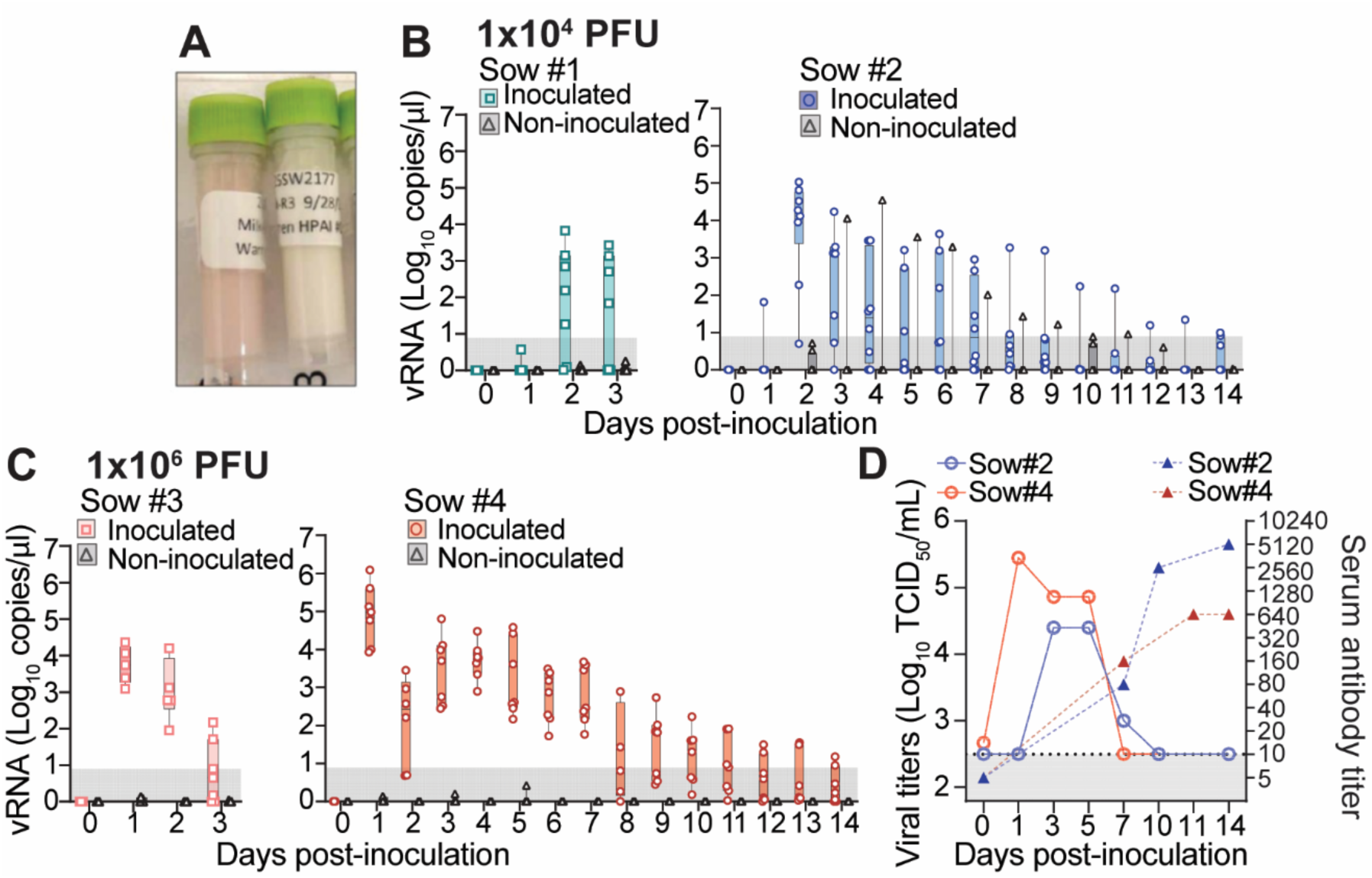
Viral shedding dynamics and seroconversion in lactating sows. Commercial lactating sows were challenged with influenza A(H5N1) clade 2.3.4.4b (D1.1 genotype) at doses of 1×10^4^ or 1×10^6^ PFU per sow, with virus equally distributed across one-half of actively lactating mammary glands. (**A**) Representative milk samples from inoculated (left) and non-inoculated (right) glands, showing changes in milk color and consistency. (**B, C**) Milk collected from inoculated and non-inoculated teats were evaluated for viral RNA (vRNA) by RT-qPCR. Data are presented as box and whisker plots of Log_10_ vRNA copy number per μL, with individual points representing milk samples from a single teat. Values plotted at 0 indicate samples with no detectable amplification signal by RT-qPCR. Shaded regions represent the limit of detection for the assay. (**D**) Milk samples were pooled from inoculated teats of sow #2 and sow #4 at the indicated days post infection (X-axis) and live virus shedding was quantified using TCID_50_ assays (left Y-axis; solid lines). Humoral responses for both sows were evaluated via live virus neutralization test (VNT) on longitudinal serum samples, expressed as reciprocal neutralizing titers (right Y-axis; dashed lines). Shaded regions represent the limits of detection for both assays.

To quantify viral shedding, milk samples collected from individual inoculated and non-inoculated teats were analyzed for viral RNA abundance by RT-qPCR. In the 1×10^4^ PFU group, nearly all samples from inoculated glands were positive by 2 dpi, with viral RNA levels ranging from 10^1^ to 10^5^ copies/μL (square and circle datapoints in **Fig. 3B**), indicating variability in virus output among individual glands. Viral RNA levels declined after one week and, in most cases, fell below the limit of detection prior to the study endpoint. In contrast, samples from non-inoculated glands were largely negative, with only a single gland testing positive across multiple days (triangle datapoints in **Fig. 3B**). The shedding kinetics in this gland were comparable to those observed in inoculated glands, suggesting that viral spread to adjacent or non-inoculated mammary tissue can occur, albeit infrequently. In the 1×10^6^ PFU group, viral RNA was detected in milk from all inoculated glands, with peak levels observed at 1 dpi. Compared to the low-dose group, viral RNA levels were more uniform across individual glands and remained above the limit of detection for a longer duration. No viral RNA was detected in milk from non-inoculated glands in this group (**Fig. 3C**). Live virus titrations performed on pooled milk samples from inoculated glands of sow #2 and sow #4 confirmed the presence of infectious virus in milk, with shedding kinetics that closely mirrored viral RNA levels (left axis in **Fig. 3D**). Serum virus neutralization assays demonstrated a lack of baseline neutralizing activity against the challenge virus, followed by the development of H5N1-specific humoral responses after intramammary challenge (right axis in **Fig. 3D**). Collectively, these data demonstrate that, despite the absence of overt clinical disease (**Fig. 2**), lactating sows with pre-existing IAV immunity can be productively infected with influenza A(H5N1) following intramammary exposure, shed both viral RNA and infectious virus in milk, and develop virus-specific neutralizing antibody responses.

### Tissue viral RNA and antigen distribution following intramammary inoculation

Sows from both dose groups were euthanized at 3 and 14 dpi to assess tissue-level viral RNA and antigen distribution. At 3 dpi, viral RNA was readily detectable in glandular tissue and teat cistern compartments of inoculated glands, irrespective of dose (**Fig. 4A, 4C; Supplementary Table 3**). In contrast, viral RNA was largely undetectable in non-inoculated mammary tissues, with one exception from the dose group of 1×10^4^ PFU: sow #1, left teat 2 (L2) (**Fig. 4A**). This isolated detection may reflect cross-contamination during necropsy, as milk from non-inoculated glands of this animal did not contain detectable viral RNA at the corresponding timepoint (**Fig. 3B**). By 14 dpi, viral RNA levels were at or below the limit of detection in nearly all sampled teat and gland cisterns tissues (**Fig. 4B, 4D; Supplementary Table 3**), consistent with viral clearance. Immunohistochemical labeling further confirmed these findings. In the low-dose sow, multifocal nuclear and cytoplasmic labeling for IAV nucleoprotein (NP) at 3 dpi was detected predominantly within ducts adjacent to glandular tissue. In the high-dose sow at the same timepoint, NP labeling was similarly detected within mammary gland tissue, with positive cells displaying morphology consistent with alveolar epithelial cells. In contrast, non-inoculated mammary gland tissues that were negative for viral RNA by RT-qPCR at necropsy showed no detectable NP labeling in either dose group (**Supplementary Fig. 1**). Assessment of systemic tissues revealed limited viral RNA detection in supramammary and mediastinal lymph nodes. Aside from these findings, viral RNA was not detected in other extra-mammary tissues examined, indicating a lack of systemic dissemination (**Supplementary Tables 3, 4**).

**Figure 4.**
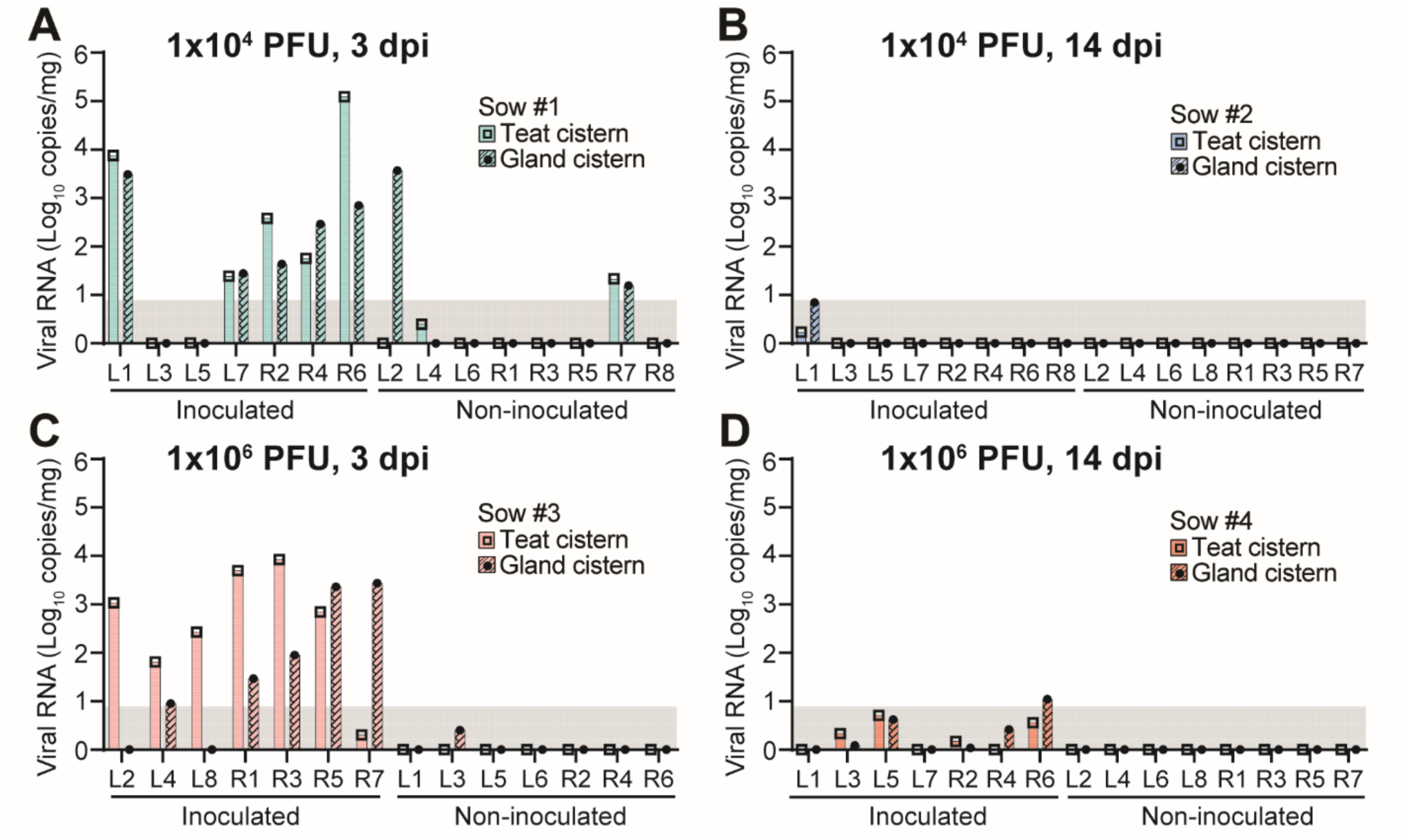
Viral RNA abundance in mammary and teat tissues. Commercial lactating sows were challenged with influenza A(H5N1) clade 2.3.4.4b (D1.1 genotype) at doses of (**A, B**) 1×10^4^ or (**C, D**) 1×10^6^ PFU per sow, with virus equally distributed across one-half of actively lactating mammary glands. At necropsy, performed at (**A, C**) 3 days post-inoculation (dpi) or (**B, D**) 14 dpi, teat cistern and gland cistern tissues were collected from both inoculated and non-inoculated mammary glands, and viral RNA abundance was evaluated by RT-qPCR. Individual mammary glands were labeled sequentially as L1–L7 (or L8) and R1–R7 (or R8), corresponding to glands on the left and right sides, respectively. Values plotted at 0 indicate samples with no detectable amplification signal by RT-qPCR. Shaded regions represent the limit of detection for the assay.

### Tissue histopathology following intramammary inoculation

Histological evaluation of mammary tissues revealed dose-dependent pathological changes following intramammary inoculation. Non-inoculated glands from low dose animals (1×10^4^ PFU; sows 1 and 2 at 3 and 14 dpi, respectively) were within normal histological limits (**Fig. 5 A-B, E-F**). In contrast, non-inoculated glands from high-dose animals (1×10^6^ PFU; sows 3 and 4) exhibited mild to moderate inflammatory changes, including infiltration of lymphocytes and plasma cells and epithelial disruption (e.g., hypertrophy, hyperplasia, and flattening) (**Fig. 5 I-J, M-N**). These findings suggest that higher inoculation doses may induce localized immune responses extending beyond directly inoculated glands.

**Figure 5.**
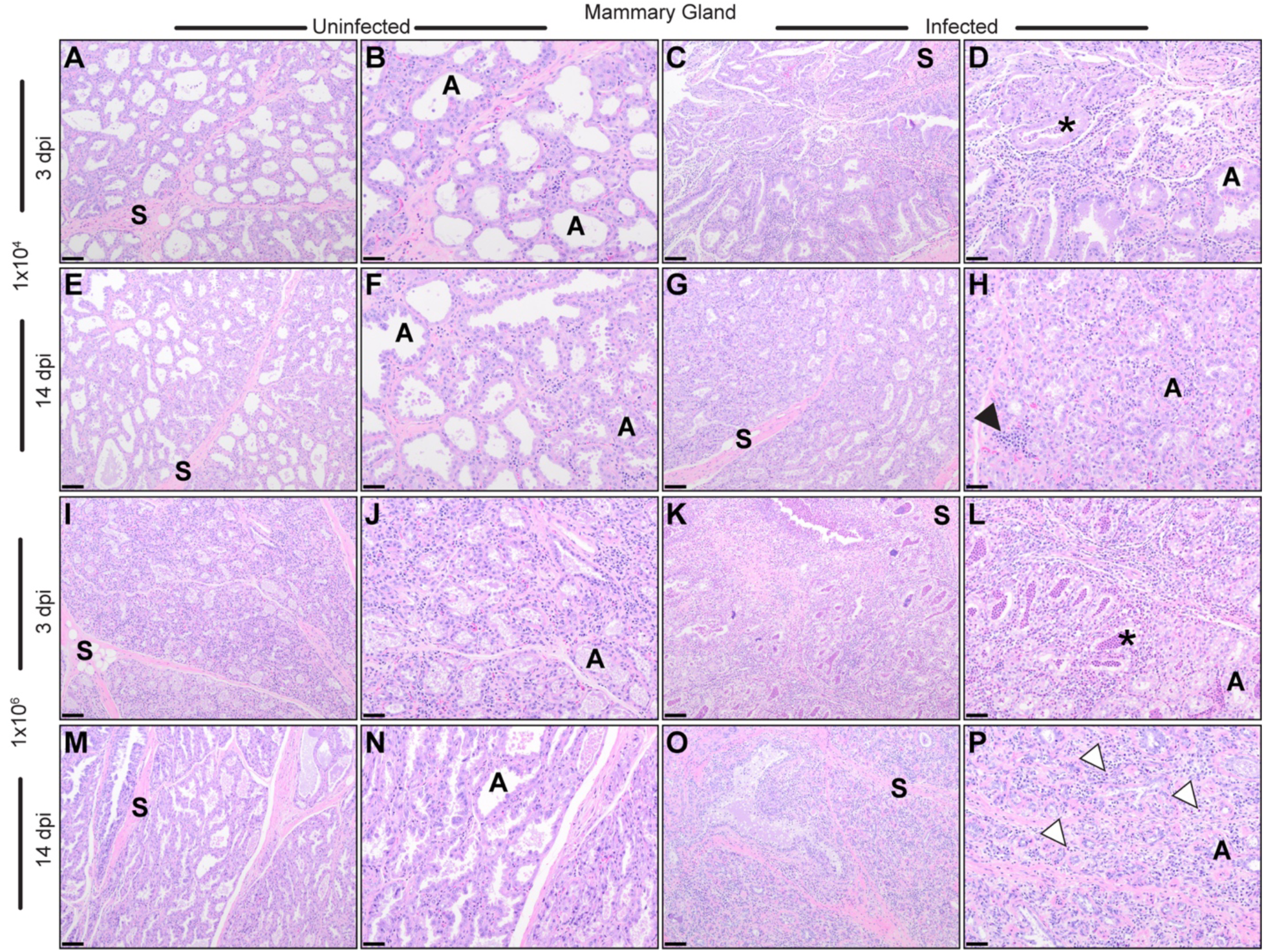
Histological examination of mammary gland tissue. Non-inoculated glands from low-dose sows at 3 and 14 dpi (sows 1 and 2 respectively, **A-B**, **E-F**) were histologically unremarkable and similar. At 3 dpi, the inoculated gland (**C-D**) had interstitial infiltrates of inflammatory cells between alveoli, which were sometimes filled with necrotic and inflammatory debris (asterisk). At 14 dpi (**G-H**), the inoculated gland showed signs of tissue remodeling and repair including regions of smaller alveoli lined by plump basophilic epithelial cells. Clusters of residual inflammatory cells, including numerous plasma cells (arrowhead), were present in the interstitium. In contrast, non-inoculated glands from high-dose sows at 3 and 14 dpi (sows 3 and 4 respectively, **I-J**, **M-N**) had lesions consistent with interstitial infiltrates of inflammatory cells at cellular debris in alveoli acutely, and evidence of epithelial injury, including variable hypertrophy/hyperplasia/attenuation acutely and chronically. Inoculated glands from the same high-dose sow at 3 dpi (**K-L**) had widespread necrosis, sheets of large numbers of neutrophils (asterisk), loss of alveoli, and interstitial infiltration by numerous inflammatory cells. By 14 dpi (**O-P**), small and compact alveoli lined by basophilic epithelium were surrounded by abundant mononuclear inflammatory cells mixed with numerous eosinophils (white arrowheads) and prominent active fibroblasts. S = collagenous interlobular septa; A = alveolus. 100x total magnification with 100 micrometer scale bar: **A, C, E, G, I, K, M, O**. 200x total magnification with 50 micrometer scale bar: **B, D, F, H, J, L, N, P**.

In inoculated glands, lesions were evident by 3 dpi and were more severe in the high-dose group. Both low- and high-dose animals exhibited epithelial necrosis and inflammatory infiltrates composed primarily of neutrophils and macrophages within glandular and teat tissues (**Fig. 5 C-D, K-L**; **Supplementary Fig. 2**). However, lesions in the high-dose sow were markedly more severe, with extensive neutrophilic inflammation, necrotic debris, and swelling of the alveolar epithelium (compare sow 3 in **Fig. 5 K-L** to sow 1 in **Fig. 5 C-D**). By 14 dpi, lesions in the low-dose animal were consistent with resolving inflammation and tissue repair (**Fig. 5 G-H**). In contrast, the high-dose animal exhibited more advanced pathology, including loss of alveolar architecture, interstitial fibrosis, persistent inflammatory infiltrates, and residual necrotic or neutrophilic debris (**Fig. 5 O-P**).

Teat lesions generally paralleled those observed in corresponding mammary glands, with severe acute inflammation at 3 dpi and more reparative or tissue remodeling changes by 14 dpi (**Supplementary Fig. 2**). Evaluation of lymphoid tissues revealed minimal involvement, with no lesions observed in mediastinal lymph nodes and only mild to moderate changes in mammary lymph nodes, most prominently in high dose animals (**Supplementary Fig. 3**). Collectively, these findings demonstrate that intramammary infection with influenza A(H5N1) results in localized mammary gland pathology that varies by dose and time post-infection, progressing from acute inflammatory injury at early timepoints to resolving inflammation or tissue remodeling at later timepoints.

### Virus transmission to piglets

To evaluate potential sow-to-piglet transmission following intramammary infection, we longitudinally collected swab samples from both lactating sows and their piglets after challenge with bovine-derived influenza A(H5N1) D1.1 virus. Nasal and rectal swabs were collected daily from sows to assess respiratory and gastrointestinal shedding, respectively, and viral RNA abundance was measured by RT-qPCR. Viral RNA was not detected in these samples, suggesting that intramammary inoculation did not result in detectable respiratory or gastrointestinal shedding in lactating sows (**Supplementary Figure 4**).

Piglets were reunited with their respective sows at 24 hours post-challenge to evaluate potential nursing-associated exposure and transmission. Oral, nasal, and rectal swabs were collected from piglets every other day throughout the study period and analyzed for viral RNA by RT-qPCR. Viral RNA was sporadically detected in oral swab samples from piglets, primarily between 2 and 6 dpi (**Fig. 6A**). The inconsistent and transient detection suggests the absence of sustained viral replication in the oral cavity. Instead, the presence of viral RNA in oral swabs is likely attributable to residual contamination associated with suckling behavior, as supported by the similarity in viral RNA abundance patterns between oral swabs and milk samples. In contrast, viral RNA was not consistently detected in nasal (**Fig. 6B**) or rectal swabs (**Fig. 6C**) from piglets, indicating a lack of measurable respiratory or gastrointestinal shedding.

**Figure 6.**
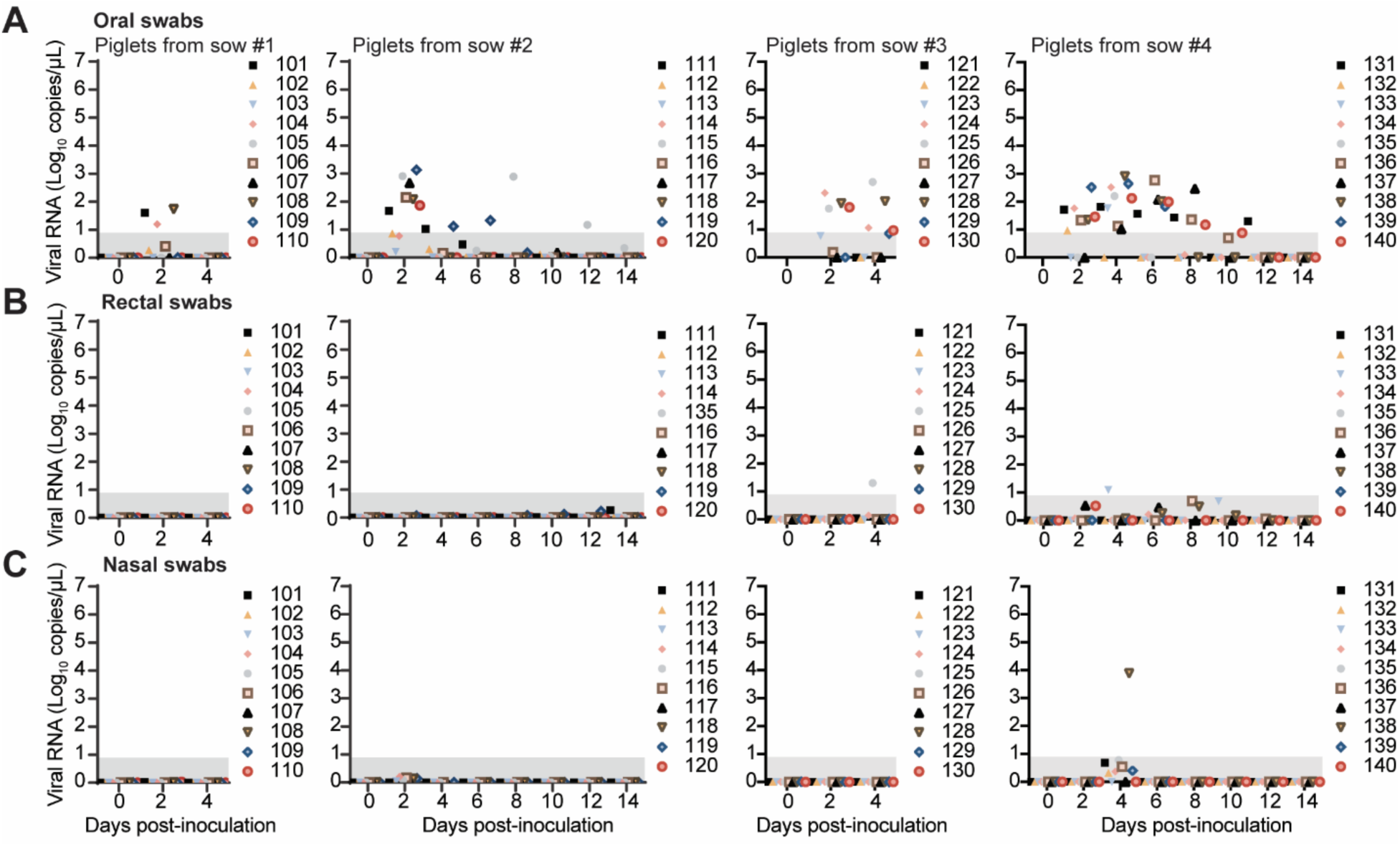
Virus transmission to piglets. 24 hours following virus inoculation into lactating sows, piglets were returned to the farrowing crate and allowed to suckle. (**A**) Oral, (**B**) rectal, and (**C**) nasal swabs were taken every other day from piglets, and viral RNA abundance was measured using RT-qPCR. Each data point represents Log_10_ copies of viral RNA per μL from an individual piglet. Values plotted at 0 indicate samples with no detectable amplification signal by RT-qPCR. Shaded regions represent the limit of detection for the assay.

## DISCUSSION

The unprecedented spillover of influenza A(H5N1) virus into U.S. dairy cattle has raised concerns regarding the risk to commercial swine populations. While other groups have interrogated respiratory transmission pathways^25,26^, none to-date have determined whether the lacatating sow mammary gland is permissive to influenza A(H5N1) virus infection. Virus transmission from cow-to-cow is suspected to occur through contaminated milk during milking procedures^17,36,41^. Although lactating sows are not mechanically milked like dairy cattle, piglets suckle continuously for 3–4 weeks throughout lactation. This prolonged close contact may provide a route for mammary exposure, whereby virus present in the oral or nasal cavity of piglets, or on the teat surface, could access the teat canal during nursing. Our findings demonstrate that, like cattle, the mammary gland of lactating swine can support productive influenza A(H5N1) virus infection and infectious virus shedding in milk (**Fig. 3**). Although direct intramammary inoculation is an artificial exposure route, this approach has been useful for defining mammary gland susceptibility in dairy cattle and allowed us to directly assess whether the swine mammary gland can support virus replication. To better approximate natural transmission dynamics, future studies should evaluate whether respiratory or oral exposure of suckling piglets can result in subsequent mammary exposure and infection of lactating sows during nursing.

For this study, we selected a bovine-origin influenza A(H5N1) genotype D1.1 virus isolated during a 2025 outbreak on a Nevada dairy farm. D1.1 viruses have become an epidemiologically important genotype in North American wild birds and domestic poultry and have been associated with multiple spillovers into dairy cattle^6,30,31^. These viruses have also been associated with over a dozen human infections, including severe and fatal cases^42–46^. Notably, the strain used in this study contains the polymerase basic 2 (PB2) D701N substitution, a known mammalian adaptation marker^5,47,48^. Our finding that this bovine-origin D1.1 virus can productively infect the swine mammary gland and be shed as infectious virus in milk underscores the potential agricultural and public health threat posed by these viruses if introduced into commercial swine production systems. As an initial investigation, this study underscores the threat of a concerning emerging genotype; however, further studies involving diverse influenza A(H5N1) clade 2.3.4.4b genotypes circulating in wild birds are warranted to more comprehensively evaluate the risk to commercial swine populations.

A key feature of this study was the use of lactating sows with prior IAV vaccination histories representative of commercial swine populations, allowing us to directly and rapidly evaluate the threat of influenza A(H5N1) to the swine industry. Although endemic IAV exposure and vaccination are common in U.S. swine herds, commercial inactivated vaccines have been reported to provide limited cross-protection against antigenically distinct emerging variants^49–51^. Thus, high IAV seropositivity within herds should not be interpreted as evidence that commercial pigs are protected from H5N1 virus infection. Following intramammary exposure to bovine-origin influenza A(H5N1) genotype D1.1 virus, vaccinated lactating sows developed productive, localized mammary infection with prolonged viral shedding in milk (**Figs. 3-5**, **Supplementary Figure 1**, **Supplementary Tables 3 and 4**), yet exhibited no systemic or respiratory clinical signs (**Fig. 2**). It remains unclear whether prior immunity against endemic swine IAV prevents clinical diseases following influenza A(H5N1) infection. Heterologous protection may be provided by certain strains^52^, and could potentially be conferred to nursing piglets via maternally derived antibodies. Future studies should include immunologically naïve sows and piglets to evaluate how immune history influences mammary infection, shedding kinetics, and clinical outcomes. Here, the evidence of mammary infection in commercial sows suggests that surveillance strategies relying solely on respiratory signs or overt herd-level illness may fail to detect localized mammary infection in lactating swine.

Viral replication was largely restricted to inoculated mammary tissues, with only one non-inoculated gland showing evidence of infection (**Fig. 3B**). Compartmentalized mammary infections have similarly been reported following experimental influenza A(H5N1) virus infection in cows and sheep^37,38,40,53^. However, the anatomical context of the mammary gland differs amongst these livestock species: ruminants possess two udder halves (sheep) or four quarters (dairy cattle), whereas sows possess 12–18 anatomically distinct mammary glands with independent glandular parenchyma^54^. Although we cannot rule out direct spread between adjacent mammary glands in swine with our experimental design, intermittent oral viral RNA positivity in suckling piglets (**Fig. 6**) raises the possibility virus may be mechanically transferred between glands during nursing. Studies in other lactating mammalian models, including dairy cattle, goats, and sheep, support the plausibility of retrograde mammary exposure during suckling^53,55,56^. Thus, while productive piglet infection was not demonstrated in this study, localized mammary infection and mechanical transmission via nursing-piglets could contribute to silent dissemination under field conditions.

Collectively, this study expands the host and tissue contexts in which contemporary influenza A(H5N1) viruses may threaten agricultural production and public health. The clinically inapparent presentation in sows suggests that observation for overt disease alone may be insufficient to detect infection in swine. As a common limitation for large-animal studies conducted under BSL-3Ag containment, the small sample size (n = 4 sows) restricts our ability to evaluate animal-to-animal variability or perform robust dose-dependent comparisons. However, similar patterns of viral RNA shedding, infectious virus recovery, and mammary tissue pathology, as well as viral RNA positivity in piglets, support the conclusion that lactating sows are susceptible to influenza A(H5N1) infection following intramammary exposure. Given the role of swine in IAV evolution and reassortment, early detection of influenza A(H5N1) infection in this host population should be a priority for animal and public health preparedness.

## MATERIALS AND METHODS

### Ethics and biosafety

The virus experiments and animal study were approved and performed under The Ohio State University Institutional Biosafety Committee (IBC, Protocol #2025R00000037) and the Institutional Animal Care and Use Committee (IACUC, Protocol #2025A00000035) in compliance with the Animal Welfare Act. All animal and laboratory work were conducted in biosafety level 3 agriculture (BSL-3Ag) containment and BSL-3 laboratory in The Ohio State University Ralph Regula Plant and Animal Agricultural Research (PAAR) facility in Wooster, OH, USA except for inactivated samples that were processed at BSL-2.

### Cell culture

Madin-Darby canine kidney (MDCK; ATCC #CCL-34) epithelial cells were maintained in Eagle’s Minimum Essential Medium (EMEM; Fisher Scientific, #10–009-CV) supplemented with 10% fetal bovine serum (FBS; Millipore Sigma, #F2442–500ML) and 1% Penicillin-Streptomycin (Pen Strep; Gibco #15140163) (complete EMEM) at 37 °C in a humidified 5% CO_2_ atmosphere.

### Virus propagation and titration

#### Virus propagation

The bovine-derived influenza A(H5N1) D1.1 virus, A/bovine/Nevada/WD-210/2025 (GenBank IDs: PV520600.1–PV520607.1), was propagated on MDCK cells as follows. Cells were seeded the day prior to infection on T75 flasks at 7.5×10^6^ cells/per flask in complete EMEM. The following day, the media was removed, the cells were washed 1X with PBS, and replaced with 4 mL of virus-containing media (1:1000 dilution of A/bovine/Nevada/WD-210/2025 [stock HA titer of 320] in EMEM with 1% Pen Strep and 0.3% bovine serum albumin [BSA]). The virus exposed cells were incubated at 37 °C in 5% CO_2_ for 1 h with manual rocking every 15 min to allow virus attachment. Following incubation, the inocula were removed and replaced with influenza growth media (EMEM containing 1% Pen Strep, 0.3% BSA, and 1 μg/mL of TPCK-treated trypsin) and returned to the incubator. Cell supernatants were harvested when the cell monolayer showed >70% cytopathic effect (CPE). The supernatant was centrifuged at 1000xg for 5 min to pellet cell debris and then aliquoted into single use vials and stored at −80 °C.

#### Virus titration by plaque assay

Virus stock samples were 10-fold serial-diluted with serum free EMEM, and 800 μL of each diluted samples were added to confluent MDCK cells (1×10^6^ cells per well in 6-well plates), followed by incubation at 37 °C in 5% CO_2_ for 1 h with gentle rocking every 15 min. Inocula were then removed, and a volume of 3 mL of overlay media (final concentration 1X phenol-free EMEM [Neta Scientific, #QB-115-073-101] + L-glutamine [Life Technologies, #25030081] + Pen Strep + 1.4% Avicel [IFF Pharma Solutions, #RC-581] + 1 μg/mL TPCK trypsin) was added to each well, followed by undisturbed incubation at 37 °C for 48 h. Following incubation, the overlay media was removed, and cells were then washed twice with PBS, and then fixed with crystal violet solution (20% methanol containing 0.2% crystal violet in PBS) for 30 min. Plaques were visualized following the removal of the crystal violet solution and 2x washes with PBS. Viral titers were calculated as plaque-forming units per milliliter (PFU/mL).

### Experimental design

A total of four lactating sows (*Sus Scrofa domesticus*; cross-bred) and forty 1-week-old piglets (ten piglets born from each sow) were obtained from a commercial breeding farm in Ohio and transported to a BSL-3Ag facility. According to farm records, all sows were of fourth parity and had previously had five administrations of swine influenza vaccines targeting multiple genotypes. Vaccinations were administered every 20 weeks, alternating between prime and boost doses. The prime vaccine targeted the H1N1 1A.1.1, H1N1 1A.3.3.3, H3N2 3.1990.4.a, and H3N2 3.2010.1 strains, while the boost vaccine targeted the H1N1 1A.1.1, H1N1 1A.3.3.2, H1N2 1B.2.1, H3N2 3.1990.4.a, and H3N2 3.2010.1 strains. All sows received a prime dose as their final vaccination approximately 20 weeks prior to the initiation of this study. To standardize litter size for experimental evaluation, piglet numbers were adjusted to ten piglets per sow prior to animal shipment by cross-fostering excess piglets to other lactating sows on the source farm. Animals were housed under BSL-3Ag containment and acclimated for three days prior to experiments. Sows were provided a gestation/lactation diet fed as a gruel (Centerra; 1:2 feed-to-water mixture), with *ad libitum* access to water. Sows were randomly assigned to two dose groups (n = 2 per group) and inoculated with either 1×10^4^ or 1×10^6^ PFU of the bovine-derived influenza A(H5N1) D1.1 virus (A/bovine/Nevada/WD-210/2025) via the intramammary route. To prepare the inoculum, virus stocks were first resuspended in 5 mL of EMEM (no extra additions) and distributed into individual syringes (0.5 mL volume per inoculated gland). Following disinfection of teat skin and orifice with povidone-iodine and 70% ethanol, the inoculum was infused into teat canals of half the actively lactating mammary glands using 25-gauge sterile cannulas (Jorgensen Laboratories). Post-infusion, teats were gently massaged to facilitate distribution into gland cistern. Inoculations followed a zig-zag pattern, alternating between left and right sides, while the remaining glands were left non-inoculated. Litters of piglets were separated from sows for 24 h following inoculation to allow for establishment of intramammary infection without suckling interference. Piglets were provided with milk replacer during the period. At 24 h post-inoculation, litters were reunited with the sows to allow natural nursing and potential viral exposure.

Animals were monitored daily for clinical signs, including lethargy, anorexia, respiratory distress, gastrointestinal signs, and neurological deficits. Milk from sows were evaluated daily for changes in milk color and consistency, and feed intake was recorded twice daily. Rectal temperature was recorded twice daily for sows and once daily for piglets, and body weight was recorded once daily for piglets. Samples of blood, milk, nasal, oral, and rectal swabs were collected at indicated time points (**Fig. 1**) and stored at −80 °C for downstream virological and serological evaluation. To evaluate pathogenesis at different infection stages, one sow per dose group was humanely euthanized for necropsy at 3 dpi and the remaining sow at 14 dpi. The corresponding litters of piglets were euthanized on the day following the sow’s necropsy. At necropsy, gross lesions were assessed, and systemic tissues were collected for viral detection and histopathological examination.

### Baseline serological assessment of sows

#### IAV nucleoprotein (NP) ELISA

Serum samples were collected from sows prior to the study to assess baseline serological status against influenza A virus (IAV). IAV nucleoprotein (NP)-specific antibodies were detected in duplicate using a commercially available epitope-blocking enzyme-linked immunosorbent assay (ELISA) kit (Avian Influenza Virus MultiS-Screen Antibody Test Kit, IDEXX, #99-12119), according to the manufacturer’s instructions. Results were reported as sample-to-negative (S/N) ratios and measured using an Infinite^®^ 200 PRO plate reader (Tecan Life Sciences). Data acquisition and analysis were performed using the i-control™ software (Tecan Life Sciences, version 2.0.10.0). This assay has been validated for the detection of anti-influenza antibodies in swine serum, with samples exhibiting S/N ratios > 0.67 considered negative for anti-IAV NP antibodies based on optimization described by Ciacci-Zanella *et al*^57^.

#### Hemagglutination inhibition (HAI) assay

Serum samples were treated with receptor-destroying enzyme (RDE [II], Denka Seiken, #370013) and incubated at 37 °C for 18–20 h, followed by heat inactivation at 56 °C for 30 min. Treated sera were serially 2-fold diluted and incubated with 4 hemagglutination units (HAU) of virus for 1 h at room temperature. Hemagglutination was subsequently assessed using 0.5% turkey red blood cells, and HAI titers were determined in duplicate as the reciprocal of the highest serum dilution that completely inhibited hemagglutination. Viruses selected for HAI analysis represented predominant endemic swine IAV clades circulating in swine populations, including A/swine/Ohio/18TOSU4522/2018 (H1N2; 1A.1.1 clade), A/swine/Indiana/15TOSU1184/2015 (H1N1; 1A.3.3.3 clade), A/swine/Ohio/16TOSU8308/2016 (H1N2; 1B.2.2 clade), A/swine/Ohio/35/2017 (H1N2; 1B.2.1 clade), A/Ohio/28/2016 (H3N2; 3.2010.1 clade), and A/Minnesota/11/2010 (H3N2; 3.1990.4.a clade). Clade classification and nomenclature were assigned based on the ISU *FLU*ture tool^58^ at https://influenza.cvm.iastate.edu/clades.php. To evaluate H5-specific antibody responses, an attenuated reverse genetics-derived rg-A/American wigeon/South Carolina/22-000345-001/2021 (H5N1) (HA modified, NA) + PR8 [R] (6+2) virus was additionally included in the assay using 1% horse red blood cells.

### Sample collection and processing

Milk samples were collected daily from individual inoculated and non-inoculated teats of each lactating sow. Prior to milk collection, oxytocin was administered intramuscularly to sows to facilitate milk letdown. All teats were cleaned with teat-cleansing wipes (Milk Check, #64-67000BR), and milk samples were manually stripped from each teat. Gloves were changed between collection from individual teats to minimize cross-contamination risk. Nasal and rectal swabs were collected daily from sows, while oral, nasal, and rectal swabs were collected from piglets every other day throughout the study period. Blood samples were collected daily from sows through 3 dpi, with additional collections at 5, 7, 10, and 14 dpi for animals maintained to the later study endpoint. For piglets, blood samples were collected prior to exposure and at necropsy. Blood samples were collected in serum separator tubes and centrifuged at 1,200xg for 10 min to isolate serum for subsequent serological analysis. At necropsy, tissue samples were collected in 10% neutral buffered formalin for histopathology and in 800 μL of transport media (Rocky Mountain Biologicals, #VTM-CHT-01L) for virus detection. Tissue samples collected for virus detection were homogenized using a TissueLyser II (Qiagen) homogenization system at 20 Hz for 60 s followed by 30 Hz for 60 s. The homogenization cycle was repeated after reversing tube orientation within the adapter to ensure complete tissue disruption. Tissue homogenates were clarified by centrifugation at 13,800xg for 1 min, and supernatants were collected for downstream virological analysis.

### Viral RNA extraction and RT-qPCR

Viral RNA was extracted from 100 μL of each sample using the magnetic bead-based automated extraction system (MagMAX™ Viral/Pathogen Nucleic Acid Isolation Kit, Applied Biosystems, #A48310) on a KingFisher™ Flex System (Thermo Fisher Scientific), according to the manufacturer’s instructions. Viral RNA detection and quantification were performed using the VetMAX™-Gold SIV Detection Kit (Applied Biosystems) on a QuantStudio™ 5 Real-Time PCR System (Applied Biosystems). The limit of detection for the assay was defined by a cycle threshold (Ct) cut-off value of 38. Standard curves were generated for each RT-qPCR run using serial 4-fold dilutions of a kit-provided positive control with a known concentration. Viral RNA copy numbers (copies/µL) of individual samples were calculated based on linear regression of the corresponding standard curves.

### Live virus quantification by TCID_50_ assay

Infectious virus titers of sow milk samples were determined by TCID_50_ assay using MCDK cells. Samples were initially diluted 1:100 in influenza growth media (EMEM containing 1% Pen Strep, 0.3% BSA, and 1 μg/mL of TPCK-treated trypsin). Samples were subsequently serially 5-fold diluted and inoculated in quadruplicate (100 µL per well) onto confluent MDCK cells seeded at 5×10^4^ cells/well in 96-well plates. Non-inoculated wells were included on each plate as negative controls. After incubation for 72 h at 37 °C in 5% CO_2_, culture medium was removed and cells were washed with PBS. Cells were then stained for 30 min with crystal violet solution (20% methanol containing 0.2% crystal violet in PBS). After thorough PBS washes, each well was evaluated for the presence or absence of cytopathic effect (CPE), and TCID_50_ titers were calculated using the Reed–Muench method.

### Histology

Fixed tissues were routinely processed on a Leica Histocore Peloris Tissue Processor (Leica Microsystems), embedded in paraffin wax, sectioned by a certified histotechnologist, stained with Hematoxylin & Eosin (H&E) and coverslipped on a Leica HistoCore SPECTRA and CV Autostainer/Automated Coverslipper (Leica Microsystems). All histology procedures were performed by The Ohio State University Comprehensive Cancer Center Comparative Pathology & Digital Imaging Shared Resource by certified technical staff and supervised by a board-certified comparative pathologist. H&E slides were evaluated by a board-certified comparative pathologist (K. Corps).

### Immunohistochemistry

Immunohistochemistry for influenza A virus antigen was performed on formalin-fixed, paraffin-embedded tissue sections using a QA/QC-validated BioCare Medical intelliPATH FLX autostainer. Slides were routinely deparaffinized and subjected to heat-induced epitope retrieval in pre-heated citrate-based Target Retrieval Solution (Dako, S1699) for 30 min at 95 °C, followed by cooling for 15 min. Endogenous peroxidase activity was blocked using 3% hydrogen peroxide (Fisher, H325-500), and nonspecific binding was blocked with normal goat serum (Vector Laboratories, MP-7444) for 20 min. Sections were incubated for 30 min with a rabbit anti-influenza A primary antibody (GeneTex, GTX125989; 1:2000 dilution), followed by a horseradish peroxidase-conjugated anti-rabbit secondary antibody (Vector Laboratories, MP-6401) for 30 min. Immunolabeling was visualized using 3,3′-diaminobenzidine chromogen (DAB; Dako, K3468) for 2.5 min, and sections were counterstained with hematoxylin for 30 s. Washes were performed between reagent steps according to the autostainer protocol. Chromogenic immunohistochemistry was performed by a certified IHC histotechnologist and reviewed by a board-certified veterinary comparative pathologist (K. Corps).

### Live virus neutralization test

Neutralizing antibodies in serum collected from sows were quantified by virus neutralization tests (VNTs) using MDCK cells. Serum samples were heat-inactivated at 56 °C for 30 min and diluted 1:10 in influenza virus growth media (EMEM containing 0.3% BSA and 1 µg/mL TPCK-treated trypsin). Samples were subsequently serially 2-fold diluted and incubated with 70 PFU per 50 μL of influenza A(H5N1) D1.1 genotype for 1 h at room temperature. Virus–sample mixtures were then added in quadruplicate onto confluent MDCK cells (seeded at 5×10^4^ cells/well in 96-well plates) and incubated for 72 h at 37 °C in 5% CO_2_. After 72 h, cells were rinsed with PBS, placed in fixation/stain solution (PBS + 20% methanol + 0.2 % crystal violet), and manually inspected for CPE. The virus neutralization titer was defined as the highest dilution protecting at least 2 out of 4 replicate wells from CPE.

## ACKNOWLEDGEMENTS

We would like to thank PAAR facility staff (Alden Sewell, Kaitlynn Starr, Jonathan Zsoldos, and Michael Jeffers) for their support of this study. The authors gratefully thank Dr. Richard Webby (St. Jude Children’s Research Hospital) for generously providing the A/bovine/Nevada/WD-210/2025(H5N1) virus isolate, as well as the Ohio Pork Council for providing the sow housing system utilized in this study. This study was funded in part by the Swine Health Information Center (SHIC) project #25-020 and the United States Department of Agriculture (USDA) National Institute of Food and Agriculture (NIFA) under contract 2025-39601-44639 (to CJW). We thank the Comparative Pathology and Digital Imaging Shared Resource at The Ohio State University Comprehensive Cancer Center, Columbus, Ohio for the histology and immunohistochemistry, which was supported by The Ohio State University Comprehensive Cancer Center and the National Institutes of Health (P30 CA016058).

