## Supplemental Figure 1 for "Bovine-derived H5N1 influenza virus efficiently infects lactating swine via the mammary gland"

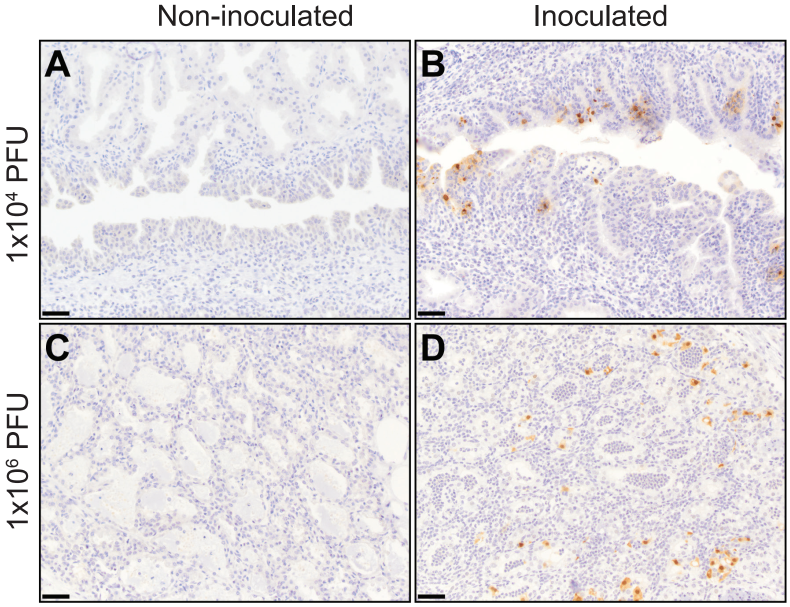


**Supplementary Figure 1. Immunohistochemical labeling of mammary tissues for low and high dose inoculated sows.** Regardless of dpi, tissues from non-inoculated glands (low dose [1x$10^4$ PFU], **A**; high dose [1x$10^6$ PFU], **C**) had no positive labeling for influenza A virus (IAV) nucleoprotein (NP). (**B**) Inoculated tissues from glands inoculated with the low dose had multifocal positive nuclear and cytoplasmic labeling for IAV NP at 3 dpi, with distribution predominantly in the ducts, particularly those immediately adjacent to the glands. (**D**) In the high dose group at 3 dpi, multifocal nuclear and cytoplasmic labeling was present in the mammary glands with positive cell morphology consistent with alveolar epithelial cells and inflammatory leukocytes. No labeling was detected in either the low-dose or high-dose group at 14 dpi (data not shown). 200x total magnification, 50 micrometer scale bars.
