## Supplemental Figure 2 for "Bovine-derived H5N1 influenza virus efficiently infects lactating swine via the mammary gland"

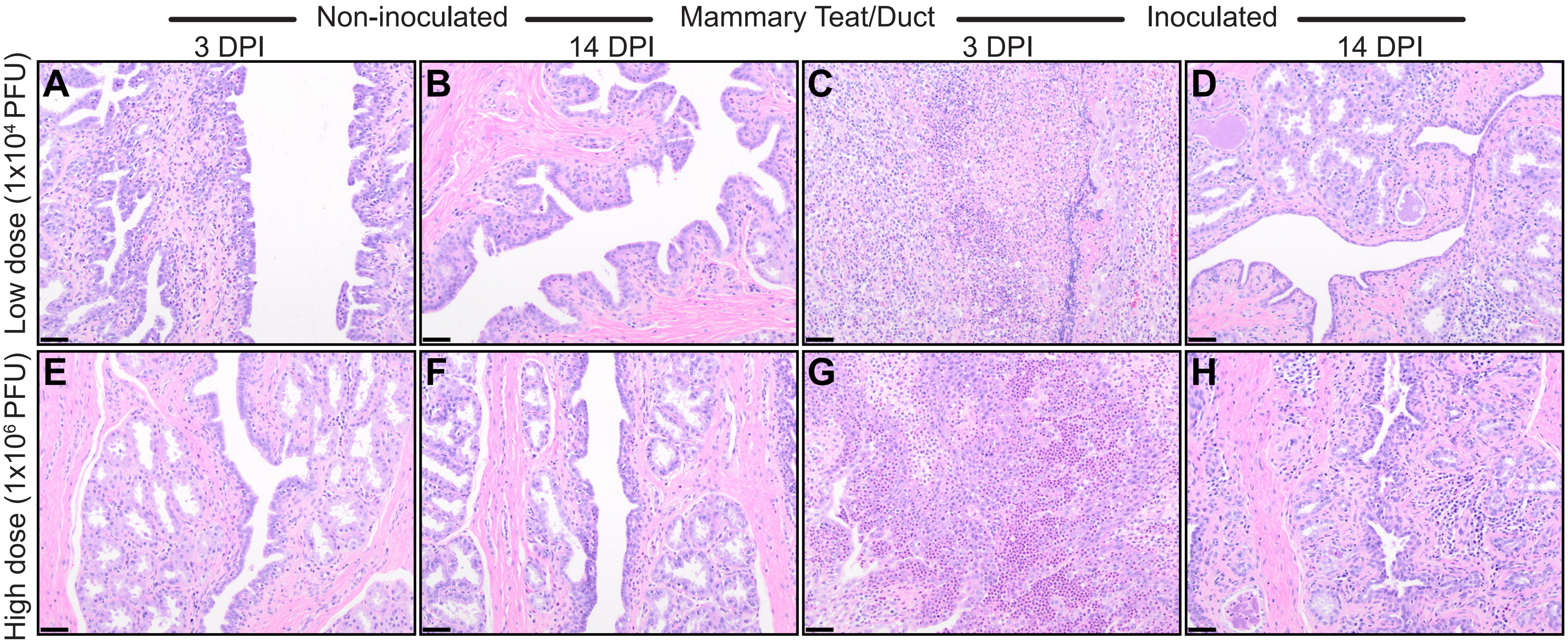


**Supplementary Figure 2. Histological examination of teat and mammary duct tissue.** Teat histopathology followed similar patterns as the associated uninoculated or inoculated glands with greater lesion severity. Regardless of dpi, teats from uninoculated glands (low-dose, **A-B**; high-dose, **E-F**) had minimal epithelial changes and variable interstitial infiltration by mixed mononuclear immune cells with variable numbers of eosinophils. Teats associated with inoculated glands had severe lesions at 3 dpi. Marked necrosis, loss of tissue architecture, and infiltration by myriad neutrophils were present in the low-dose teat (**C**), while the inoculated teat from the high-dose sow (**G**) had necrosis, degeneration of remaining epithelial cells, sheets of neutrophils similar to those seen in the gland, and infiltration by other inflammatory cells. At 14 dpi, the inoculated low-dose teat had epithelial attenuation/flattening and clusters of residual mononuclear inflammatory cells, particularly plasma cells (**D**). At the same timepoint, the teat from the high-dose sow (**H**) had epithelial hypertrophy and hyperplasia and larger numbers of residual lymphocytes and plasma cells with numerous eosinophils. 200x total magnification, 50 micrometer scale bars.
