## Supplemental Figure 3 for "Bovine-derived H5N1 influenza virus efficiently infects lactating swine via the mammary gland"

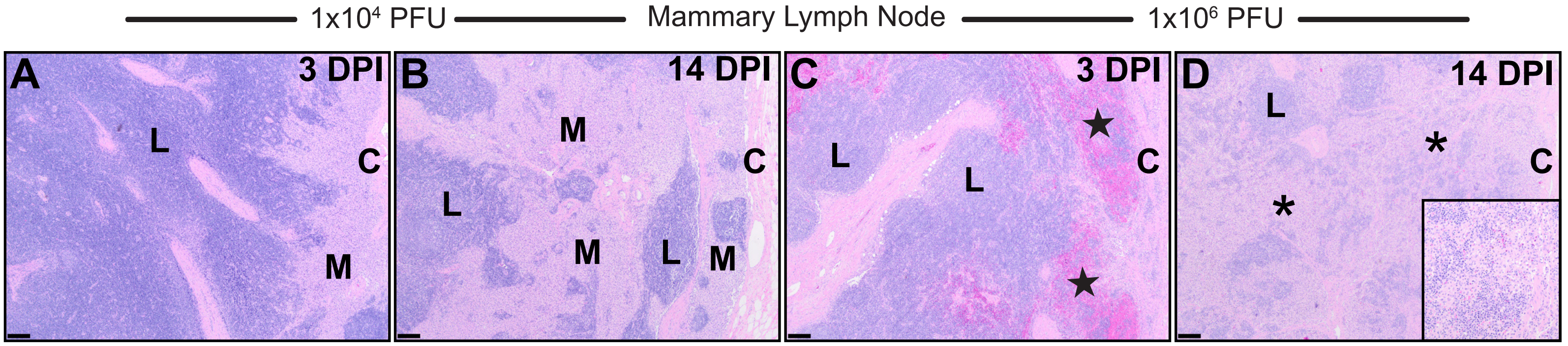


**Supplementary Figure 3. Histological examination of mammary lymph nodes following intramammary inoculation.** Mammary lymph nodes were relatively unaffected from low-dose sows at 3 and 14 dpi (**A** and **B**, respectively), with only an increase in cortical and medullary macrophages present at 14 dpi. Mammary lymph nodes from high-dose sows had more significant lesions including hemorrhage with hemosiderin-laden macrophages (stars) at 3 dpi (**C**) and decreased lymphocyte populations and infiltration by neutrophils, eosinophils, and increased macrophages (asterisks) at 14 dpi (**D**). L = lymphocytes; M = macrophages; C = capsule. 40x total magnification, 200 micrometer scale bars.
