## Supplemental Figure 4 for "Bovine-derived H5N1 influenza virus efficiently infects lactating swine via the mammary gland"

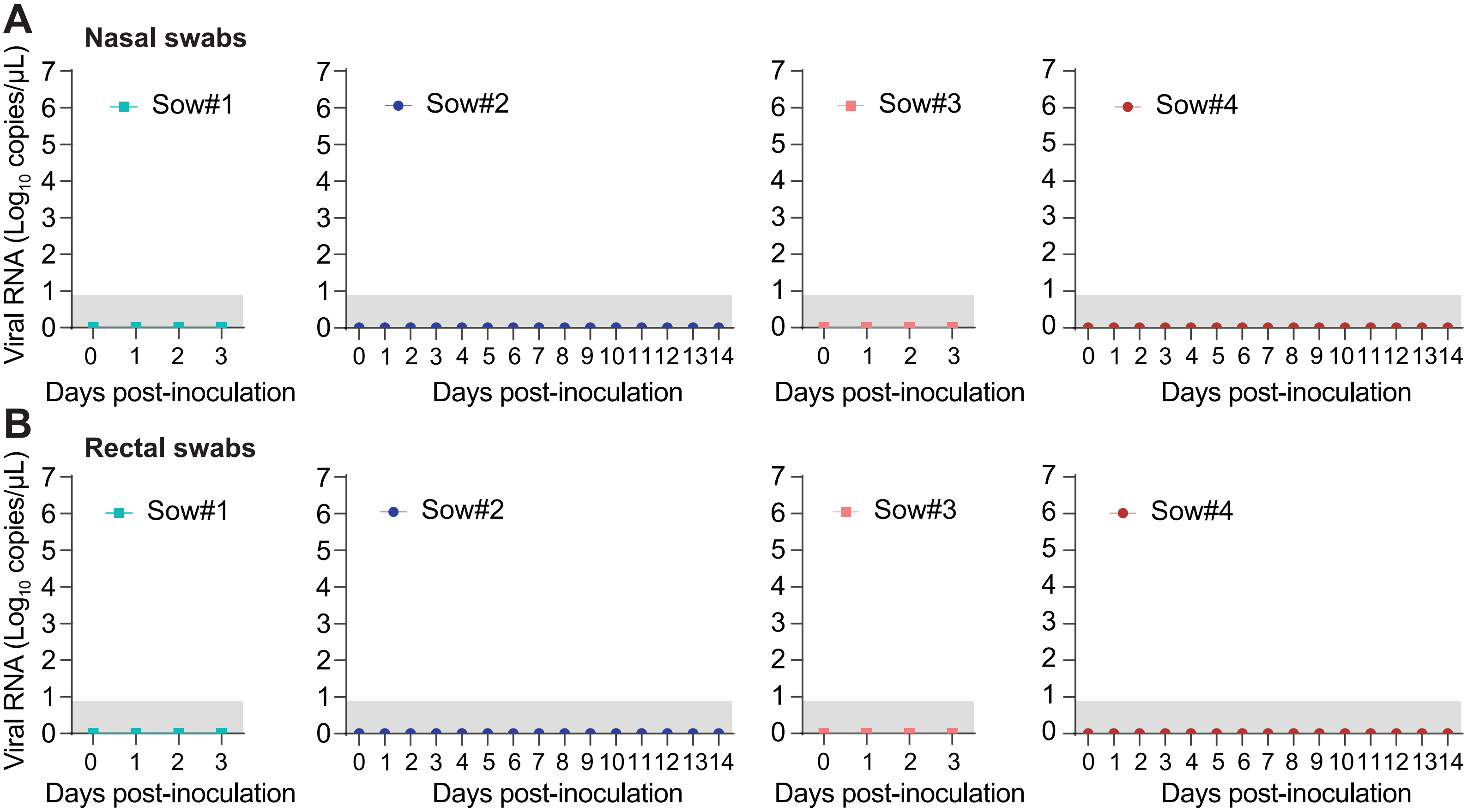


**Supplementary Figure 4. No evidence of viral RNA shedding in sow nasal and rectal swabs.** (**A**) Nasal and (**B**) rectal swabs were taken every day from sows, and viral RNA abundance was measured using RT-qPCR. Each data point represents $Log_{10}$ copies of viral RNA from the indicated sow. Values plotted at 0 indicate samples with no detectable amplification signal by RT-qPCR. Shaded regions represent the limit of detection for the assay.
