## Supplemental Table 1 for "Bovine-derived H5N1 influenza virus efficiently infects lactating swine via the mammary gland"

**Supplemental Table 1. Observations of diarrhea in piglets following exposure to influenza A(H5N1) genotype D1.1**

| [a]dpe | [b]Number of piglets with diarrhea (Piglet ID) | |
|---|---|---|
| | $1x10^4$ PFU | $1x10^6$ PFU |
| 0 | 2/20 (#103, #119) | 0/20 |
| 1 | 3/20 (#102, #115, #119) | 3/20 (#123, #125, #130) |
| 2 | 1/20 (#113) | 0/20 |
| 3 | 2/20 (#113, #119) | 0/20 |
| 4 | 2/20 (#119, #120) | 1/20 (#132) |
| 5 | 1/10 (#120) | 2/10 (#138, #140) |
| 6 | 1/10 (#112) | 3/10 (#132, #138, #140) |
| 7 | 2/10 (#112, #120) | 1/10 (#140) |
| 8 | 1/10 (#112) | 1/10 (#132) |
| 9 | 0/10 | 1/10 (#140) |
| 10 | 1/10 (#119) | 2/10 (#138, #140) |
| 11 | 1/10 (#112) | 0/10 |
| 12 | 1/10 (#119) | 0/10 |
| 13 | 1/10 (#119) | 1/10 (#138) |
| 14 | 0/10 | 1/10 (#138) |

[a]dpe: days post-exposure.
[b]Soft, yellowish, or watery feces were noted in farrowing crates or during piglet sampling. Piglets with abnormal feces were recorded individually, and the number of symptomatic piglets out of the total housed piglets is shown.
