## Supplemental Table 2 for "Bovine-derived H5N1 influenza virus efficiently infects lactating swine via the mammary gland"

**Supplemental Table 2. Observational findings of milk in lactating sows following intramammary inoculation with influenza A(H5N1) virus D1.1 genotype**

| Sow ID | [a]dpi | [b]Milk color and consistency change | | | | | | | | | | | | | |
|---|---|---|---|---|---|---|---|---|---|---|---|---|---|---|---|
| | | Inoculated mammary glands | | | | | | | Non-inoculated mammary glands | | | | | | |
| | | L1 | L3 | L5 | L7 | R2 | R4 | R6 | L2 | L4 | L6 | R1 | R3 | R5 | R7 |
| Sow#4 | 0 | - | - | - | - | - | - | - | - | NA | - | - | - | - | - |
| | 1 | NA | - | - | - | - | - | - | - | NA | - | - | - | NA | - |
| | 2 | - | - | - | - | - | - | - | - | - | - | - | - | - | - |
| | 3 | - | - | - | - | + | - | + | - | - | - | - | - | - | - |
| | 4 | + | + | + | - | + | - | + | - | - | - | - | - | - | - |
| | 5 | + | - | + | + | + | - | - | - | - | - | - | - | - | - |
| | 6 | + | - | + | + | + | - | + | - | - | - | - | - | - | - |
| | 7 | + | + | + | + | + | + | + | - | - | - | - | - | - | - |
| | 8 | + | - | + | + | + | + | + | - | - | - | - | - | - | - |
| | 9 | - | - | - | + | + | - | - | - | - | - | - | - | - | - |
| | 10 | - | - | - | - | + | - | - | - | - | - | - | - | - | - |
| | 11 | - | - | - | - | + | - | - | - | - | - | - | - | - | - |
| | 12 | - | - | - | - | + | - | - | - | - | - | - | - | - | - |
| | 13 | - | - | - | - | + | - | - | - | - | - | - | - | - | - |
| | 14 | - | - | - | - | + | - | - | - | - | - | - | - | - | - |

[a]dpi: days post-inoculation.

[b]Daily milk samples from each mammary gland were examined for changes in color and consistency. (+): Observed changes in milk appearance, including yellowish discoloration, thickened consistency, or the presence of trace blood. (-): Milk exhibiting typical, unchanged color and texture. NA (Not Applicable): Not applicable as milk yield was too low or absent to permit a visual assessment.
